# Cryptic, extensive and non-random chromosome reorganization revealed by a butterfly chromonome

**DOI:** 10.1101/233700

**Authors:** Jason Hill, Pasi Rastas, Emil A. Hornett, Ramprasad Neethiraj, Nathan Clark, Nathan Morehouse, Maria de la Paz Celorio-Mancera, Jofre Carnicer Cols, Heinrich Dircksen, Camille Meslin, Naomi Keehnen, Peter Pruisscher, Kristin Sikkink, Maria Vives, Heiko Vogel, Christer Wiklund, Alyssa Woronik, Carol L. Boggs, Sören Nylin, Christopher Wheat

## Abstract

Chromosome evolution, an important component of mico- and macroevolutionary dynamics ^1–5^, presents an enigma in the mega-diverse Lepidoptera^6^. While most species exhibit constrained chromosome evolution, with nearly identical haploid chromosome counts and chromosome-level shared gene content and collinearity among species despite more than 140 Million years of divergence^7^, a small fraction of species independently exhibit dramatic changes in chromosomal count due to extensive fission and fusion events that are facilitated by their holocentric chromosomes^7–9^. Here we address this enigma of simultaneous conservation and dynamism in chromosome evolution in our analysis of the chromonome (chromosome level assembly^10^) of the green-veined white butterfly, *Pieris napi* (Pieridae, Linnaeus, 1758). We report an unprecedented reorganization of the standard Lepidopteran chromosome structure via more than 90 fission and fusion events that are cryptic at other scales, as the haploid chromosome number is identical to related genera and gene collinearity within the large rearranged segments matches other Lepidoptera. Furthermore, these rearranged segments are significantly enriched for clusters of functionally related genes and the maintenance of ancient telomeric ends. These results suggest an unexpected role for selection in shaping chromosomal evolution when the structural constraints of monocentricq chromosomes are relaxed.

---

Butterflies and moths comprise nearly 10% of all described species^6^ and inhabit diverse niches with varied life histories, yet they exhibit a striking similarity in their genome architecture despite 140 million generations^19^ of divergence. The vast majority have a haploid chromosome number between 28 and 32^7,11,12^, and within chromosomes the gene content and order is remarkably similar among divergent species as adduced by three previous chromonomes^7,13^, BAC sequencing and chromosomal structure analyses^14,15^. Complicating this picture of conservation, haploid chromosome counts in species of Lepidoptera, as compared to all non-polyploid animals, exhibit the highest variance in number between species within a genus (n = 5 to 226^16–18^), the highest single count (n=226^9^), and polymorphism in counts that do not affect fertility in crosses^3,19^. Lepidoptera tolerate such chromosomal variation due to their holokinetic chromosomes, which facilitate the successful inheritance of novel fission or fusion fragments^20^. Thus, while Lepidoptera can avoid the deleterious consequences of large-scale bouts of chromosomal fission and fusion, in the vast majority of cases such events are only found in young clades, suggesting an evolutionary cost of chromosomal rearrangement.

The *P. napi* chromonome was generated using DNA from inbred siblings from Sweden, a genome assembly using variable fragment size libraries (180 bp to 100 kb; N50-length of 4.2 Mb and a total length of 350 Mb), and a high density linkage map generated using 275 full-sib larva, which placed 122 scaffolds into 25 linkage groups, consistent with previous karyotyping of *P. napi*^21,22^. After assessment and correction, the total chromosome level assembly was 299 Mb, comprising 85% of the total assembly size and 114% of the k-mer estimated haploid genome size, with 2943 scaffolds left unplaced (Supplementary Note 3). Subsequent annotation predicted 13,622 gene models with 9,346 functional predictions (Supplementary Note 4) and 94% of expected single copy genes (BUSCOs) were found complete (Supplementary Note 1), along with a typical mtDNA genome (Supplementary Figure 6).

The content and structure of the *P. napi* chromonome was compared with the available Lepidopteran chromonomes: the silk moth *Bombyx mori* (Bombycidae), the postman butterfly *Heliconius melpomene* (Nymphalidae), and the Glanville fritillary *Melitaea cinxia* (Nymphalidae). These latter three species exhibit gene collinearity along their chromosomes that is maintained even after chromosomal fusion and fission events, readily reflecting the history of these few events^13^ (Fig. 1). After identifying the shared single copy orthologs (SCO) among these four species’ chromonomes, we placed these results into a comparative chromosomal context. Unexpectedly, nearly every *P. napi* chromosome was uniquely reorganized on the scale of megabasepairs in what appeared to be the result of a massive bout of fission and subsequent fusion events, including the Z chromosome (Fig. 1a). A detailed comparison of the size and number of these rearrangements was then made between *P. napi* and *B. mori*, as the latter has a high quality chromonome and a haploid chromosome count (n=28) closest to the Lepidopteran mode of n=31. Using the shared SCOs, 99 well defined blocks of collinear gene order (hereafter referred to as “collinear blocks”) were identified, with each collinear block having an average of 69 SCOs. Each *P. napi* chromosome contained an average of 3.96 (SD = 1.67) collinear blocks, which derived from an average of 3.5 different *B. mori* chromosomes. In *P. napi*, the average collinear block length was 2.82 Mb (SD = 1.97 Mb) and contained 264 genes in our annotation (SD = 219).

**Figure 1:**
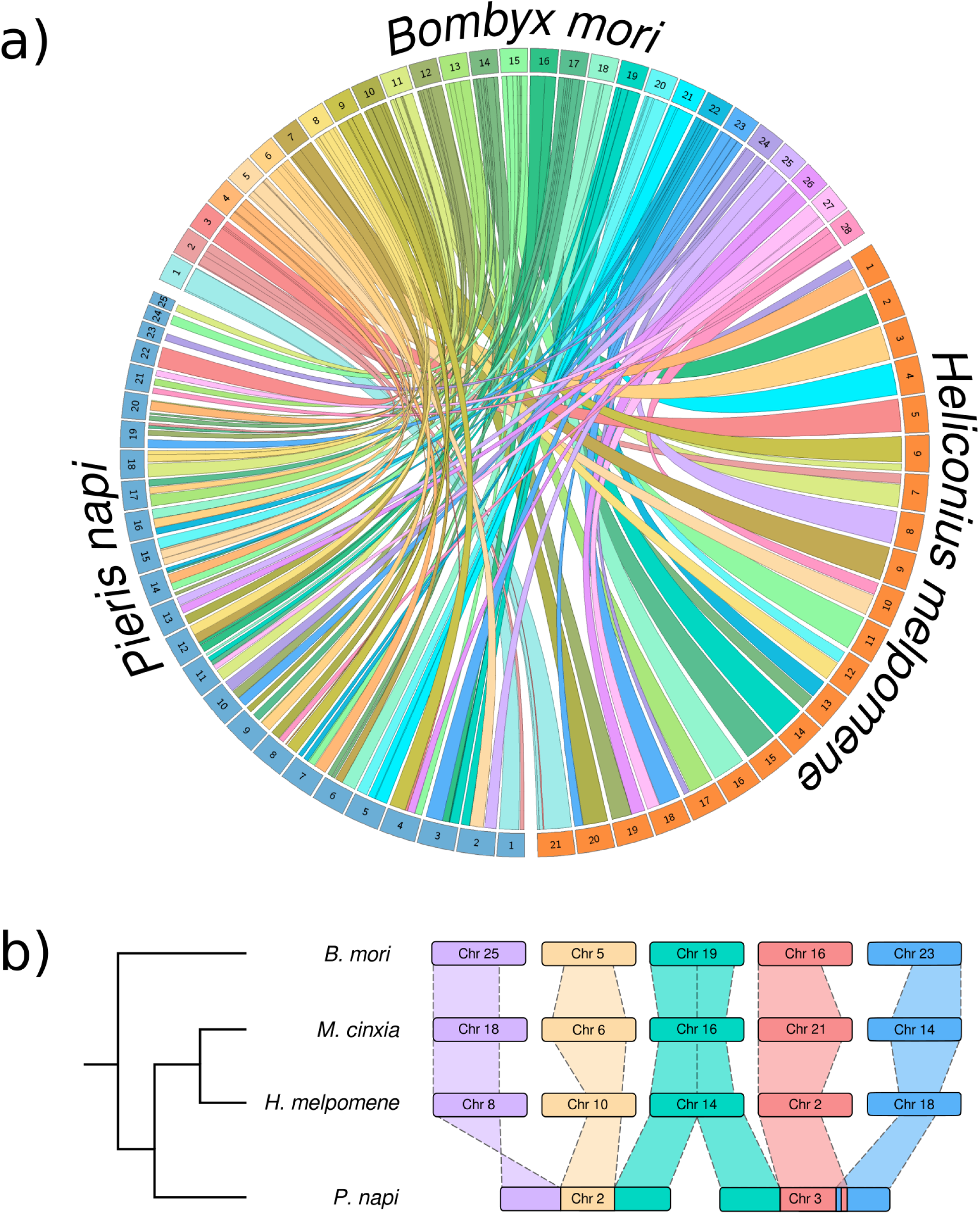
Chromosomal mapping of the moth *Bombyx mori* (Bombycoidea) to the butterflies *Pieris napi* (Pieridae), *Heliconius melpomene* and *Melitaea cinxia* (Nymphalidae)^24^. **a)** Single copy orthologs (SCOs) connecting *B. mori* to *P. napi* (n=2354) and to *H. melpomene* (n=2771). The Z chromosome is chromosome 1 in *B. mori* and *P. napi* and 21 in *H. melpomene*. Links between orthologs originate from *B. mori* and are colored by their chromosome of origin, then extend to *P. napi* chromosomes (colored blue) and *H. melpomene* chromosomes (colored orange). Links are clustered into blocks of synteny, forming colored ribbons that represent a contiguous block of genes spanning a region in both species. Chromosomes 2-25 in *P. napi* are ordered in size from largest to smallest. **b)** SCOs between the two largest autosomes of *P. napi* and the other Lepidoptera highlight the former‘s novel and derived chromosomal fusion and fission events. Here, band width for each species is proportional to the length of the inferred chromosomal region of orthology, although the individual chromosomes are not to scale.

This novel chromosomal reorganization was validated using four complementary but independent approaches to assess the scaffold joins used to make the chromosomal level assembly. First, we generated a second linkage map for *P. napi* and used this to confirmed the 25 linkage groups, our assembly scaffolds and their placement within chromosomes (Fig. 2b,c; Supplementary Fig. 2). Second, since mate-pair (MP) reads spanning the scaffold joins that were indicated by the first linkage map provide an independent assessment of validity of those joins, we quantified the number of MP reads spanning each base pair position along chromosomes, revealing consistent support for the scaffold joins within chromosomes (Fig. 2a; Supplementary Fig. 2, Note 7). Third, we aligned the scaffolds of a recent high quality genome of *P. rapae*^23^ to our *P. napi* chromonome, identifying *P. rapae* scaffolds that spanned the scaffold joins within *P. napi*, finding support for 71 of the 97 joins (Supplementary Fig. 5). Fourth, since the *P. napi* scaffold joins are not associated with *B. mori* collinear blocks of SCOs, we identified boundaries of *B. mori* collinear blocks that spanned scaffold joins within *P. napi* chromosomes, finding support for 62 of the 97 scaffold joins in *P. napi* and none against them (Fig. 2d; Supplementary Fig. 2, Note 8,9). Thus, the *Pieris napi* chromonome is well supported.

**Figure 2:**
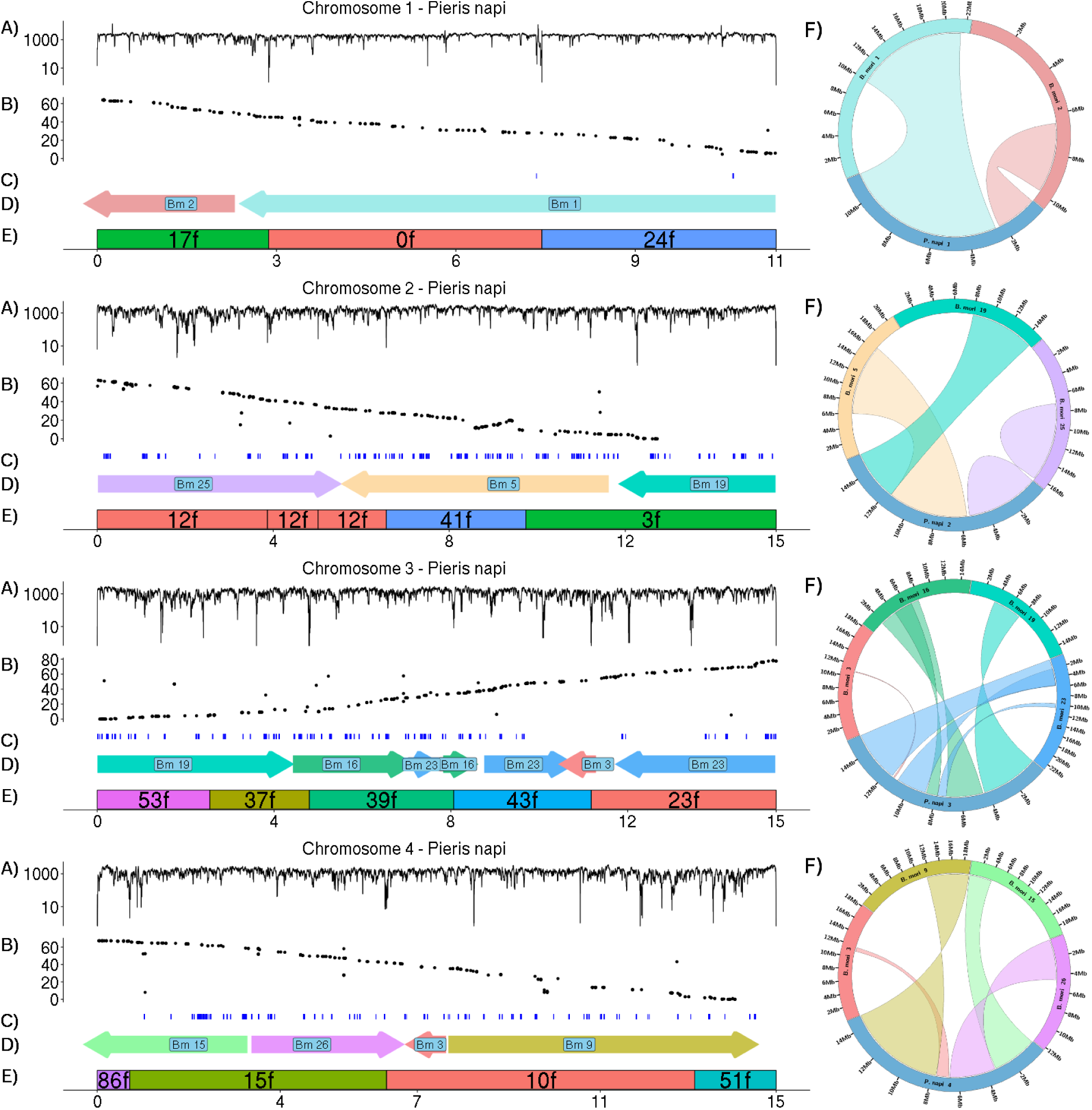
Validation of the largest four *P. napi* chromosomes. Within each, **a)** mate pair spanning depth is shown across each chromosome, summed from the 3kb, 7kb, and 40kb libraries (genome averaged = 1356). Of the scaffold join positions 74 of 97 were spanned by > 50 properly paired reads (mean = 117.8, S.D. = 298.7), while the remaining 23 scaffold joins had 0 mate pair spans. **b)** black dots represent RAD-seq linkage markers and their recombination distance along chromosomes from the first linkage map **c)** Results from the second linkage map of maternally inherited markers (RNA-seq and whole genome data), where all markers within a chromosome are completely linked due to suppressed recombination in females (i.e. recombination distance is not shown on Y axis). **d)** *B. mori* collinear blocks, colored and labeled by their chromosomal origin, along with orientation by arrow, as in Fig. 1a. **e)** *P. napi* scaffolds comprising each chromosome, labeled to indicate scaffold number and orientation. **f)** To the right of each ***P. napi*** chromosome is a circos plot showing the location and orientation of the collinear blocks from each *B. mori* donor chromosome that comprise a given *P. napi* chromosome, colored as in Fig. 1a. A twist in the ribbon indicates a reversal of the 5‘ to 3‘ orientation of the *B. mori* relative to the *P. napi* chromsomes. Ribbon width on the *P. napi* chromosome is relative to the size of the collinear block. Remaining chromosomes shown in Supplementary Fig. 2.

We next compared the *P. napi* chromosomal structure to the additional genomes available in the Lepidoptera (n=20). Like most eukaryotic genomes, most Lepidopteran genome assemblies are not chromonomes, complicating comparative assessments of chromosomal structure (Supplemental Figure 6). To overcome this limitation, we queried each scaffold of each of these genomes for SCO’s that were shared with *B. mori* and *P. napi*, and then quantified whether the scaffold supported the chromosomal structure of *B. mori* or *P. napi*. While all non-*Pieris* species have more scaffolds supporting *B.mori*-like chromosomes, two independent assemblies of the congener *P. rapae* exhibited more *P. napi*-like scaffolds (Fig 3a). Finally, we assessed the evolutionary history of haploid chromosome count in the family Pieridae, by integrating for the first time nearly 200 species level observations with a temporally calibrated phylogeny. The haploid chromosomal count of *P. napi* and *P. rapae* is essentially identical to all but one species in their clade going back to a common ancestor > 30 million years divergent^24^ (Fig 3b). Thus, the chromosomal structure of the *Pieris* lineage is novel, morphologically cryptic, and possibly old.

**Figure 3:**
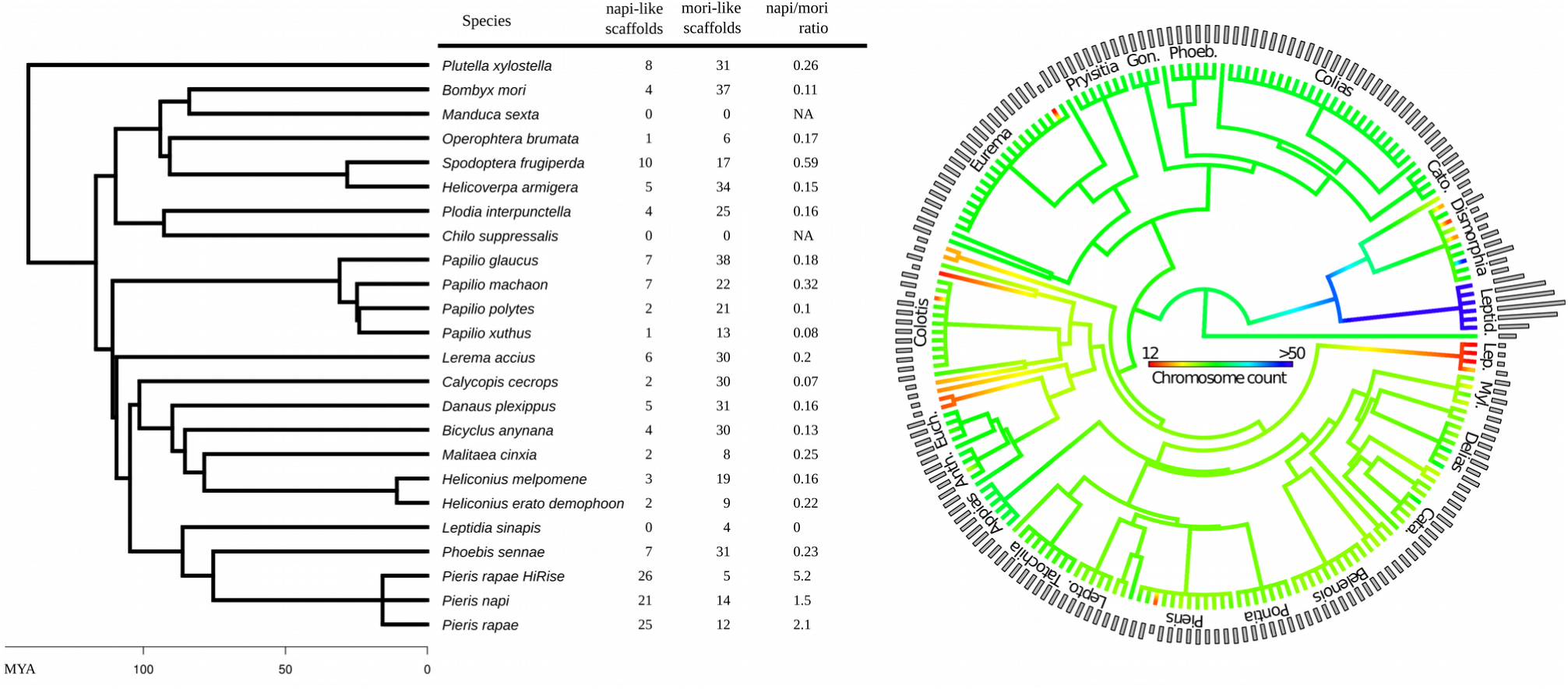
Comparative chromosomal evolution across Lepidoptera, with a focus upon the family Pieridae. a) A chronogram of currently available Lepidopteran genomes (n=24) with estimates of their chromosomal similarity relative to *B. mori* vs. *P. napi*, with time in million years ago (MYA). For each scaffold of each species, SCOs were determined with *B. mori* and *P. napi*, then we quantified the number of times a scaffold contained SCOs derived from two separate chromosomes of *B. mori*, but from a single *P. napi* chromosome (napi-like scaffold), or vice versa (mori-like scaffold)(see Supplemental Note for more details). **b)** Ancestral state reconstruction of the chromosomal fusion and fission events across a chronogram of Pieridae (n=201 species). As only a time calibrated genus-level phylogeny exists for Pieridae, all genera with > 1 species were set to an arbitrary polytomy at 5 MYA, while deeper branches reflect calibrated nodes. The haploid chromosomal count of tips (histogram) and interior branches (color coding) are indicated, with the outgroup set to n=31 reflecting the butterfly chromosomal mode. Genus names are indicated for the larger clades (all tips labels shown in Supplemental Material).

To further assess this novel chromosomal organization, we investigated the ordering and content of the collinear blocks that constitute the *P. napi* chromosomes. First, we tested whether *P. napi* maintained the traditional telomeres of Lepidoptera despite its extensive chromosomal reshuffling (Fig. 4a), finding evidence of significantly more telomeres in common between *B. mori* and *P. napi* than random chance (P < 0.01, two tailed t-test; Fig. 4b). We also identified a significant enrichment for SCOs in *B. mori* and *P. napi* located at roughly similar distance from the end of their respective chromosomes (Fig. 4c). Both of these findings are consistent with the ongoing use of ancient telomeric ends. Second, since gene order is non-random across diverse eurkaryotes^5^, we tested whether there was a gene set functional enrichment within the observed collinear blocks by investigating the full set of annotated *P. napi* genes within them. We found that 57 of the 99 collinear blocks in the *P. napi* genome contained at least three genes with a shared gene ontology (GO) term that was significantly less frequent in the rest of the genome (P < 0.01, Fishers Exact Test). We then tested whether the observed enrichment in the collinear blocks of *P. napi* was greater than expected by randomly assigning the genome into similarly sized blocks. The mean number of GO enriched fragments in each of 10,000 simulated genomes was 38.8 (variance of 46.6 and maximum of 52), which was significantly lower than the observed (P < 0.0001)(Supplementary Fig. 3). Thus, we find evidence of selection favoring the modal Lepidopteran chromosomal structure, in both the retention of ancient teleomeric ends and gene order.

**Figure 4:**
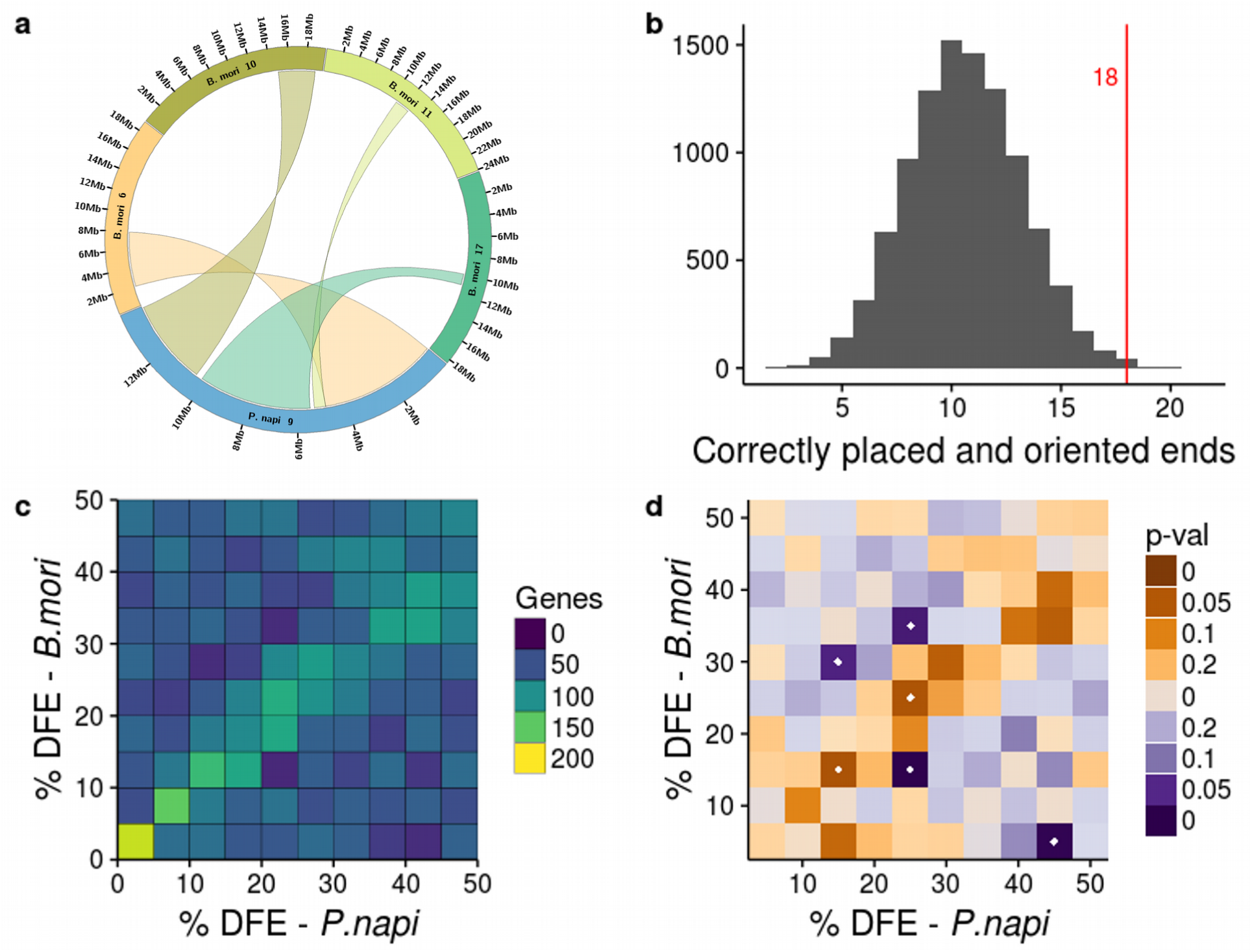
Comparison of gene content and chromosomal location of collinear blocks between *P. napi* and *B. mori* in observed and randomly permuted genomes. **a)** Observed pattern of conserved collinear block location within *P. napi* Chromosome 9, wherein telomere facing and interior origins of the syntenic blocks are conserved between species despite their reshuffling. **b)** Comparisons of the number of collinear blocks that are terminal in both *P. napi* and *B. mori* chromosomes (n=18, red line), compared to 10,000 simulated genomes (histogram: average 10.7, std dev= 6.8). **c)** Percentage distance from the end (DFE) of SCOs in a *P. napi* vs. *B. mori* chromosomes. Counts binned on the color axis. **d)** Comparison between the observed DFE distribution and the expected distribution generated from 10,000 genomes of 25 chromosomes constructed from the random fusion of the observed collinear blocks. Bins in which more genes occur in the observed genomes than the expected distribution are in orange, less genes in blue, P < 0.05 in either direction are denoted by a white dot. SCO spatial distribution was significantly higher than expected along the diagonal (two bins with p < 0.05), while significantly lower than expected off the diagonal (four bins with p < 0.05).

Repeats are known to affect chromosomal reorganization. To assess this, we surveyed the distribution of different repeat element classes across the genome, looking for enrichment of specific categories near the borders of collinear blocks. While Class 1 transposons were found to be at higher density near the ends of chromosomes relative to the distribution internally (Supplementary fig. 4), no repeat elements were enriched relative to the position of syntenic block regions. We therefore investigated whether any repeat element classes had expanded within *Pieris* compared to other sequenced genomes by assessing the distribution of repeat element classes and genome size among sequenced Lepidoptera genomes. Consistent with previous findings^25^, we observe strong relationship between genome size and repetitive element content in *Pieris* species, though no element classes appear enriched within this lineage (Supplemental Fig. 7). Thus, while repetitive elements such as transposable elements are likely to have been involved in the reshuffling, our inability to find evidence of this suggests these events may be old and their signal decayed.

The novel chromosomal organization of *P. napi* and *P. rapae* implies their ancestral lineage underwent a rampant series of fission, then fusion events. Although such chromosomal rearrangements (CRs) are well documented in Lepidoptera^8,9,11,12,15^, and the Pieridae in particular^3,15^, our evidence for a return to the near the modal haploid number of Lepidoptera after a bout of runaway fissions is unexpected. Moreover, this fragmentation and reconciliation does not appear random, as we detect the maintenance of functional gene clusters, a bias toward the use of ancient telomeric ends, and a convergence upon a common haploid count, suggesting there was a potential fitness advantage of all three. While it is tempting to envision nearly neutral selection pressures acting upon these aspects of chromosomal structure varying over time due to changes in effective population size, additional chromonomes and modeling are necessary before such scenarios can be substantiated.

## Online Content

Methods, along with any additional Extended Data display items and Source Data, are available in the online version of the paper; references unique to these sections appear only in the online paper.

**Supplementary Information** is available in the online version of the paper.

## Acknowledgements

We would like to acknowledge support from Science for Life Laboratory, the National Genomics Infrastructure, NGI, and Uppmax for providing assistance in massive parallel sequencing and computational infrastructure. We also thank SNPsaurs for support in nextRAD sequencing for linkage map construction. Funding was provided by the Swedish Research Council (VR grant numbers 2012-3715 (SN), 2010-5341 (SN), 621-2012-4001(CWW), Academy of Finland (grant number 131155 (CWW)), and the Knut and Alice Wallenberg Foundation (grant number 2012.0058).

## Author Contributions

C.W.W., S.N., J.H. designed the project. C.W. collected and inbred the material. J.H. lead the genome assembly. J.H, C.W.W., were responsible for the bioinformatic analysis of the genome. P.R. lead the linkage mapping, with help from J.H. Manual annotations were performed by R.N., N.C., N.M., M.P.C.M, J.C.C., H.D., C.M., N.K., P.P., K.S., M.V., H.V., A.W, C.B, S.N. J.H. and C.W.W. wrote the manuscript with input from other authors. All authors approved the manuscript before submission.

## Author Information

Data has been deposited on the European Nucleotide Archive, accession # PRJEB24862. Reprints and permissions information is available at www.nature.com/reprints. The authors declare no competing financial interests. Readers are welcome to comment on the online version of the paper. Publisher‘s note: Springer Nature remains neutral with regard to jurisdictional claims in published maps and institutional affiliations. Correspondence and requests for materials should be addressed to J.H and / or C.W.W.

